# The *cis*-regulatory codes of response to combined heat and drought stress in *Arabidopsis thaliana*

**DOI:** 10.1101/2020.02.28.969261

**Authors:** Christina B. Azodi, John P. Lloyd, Shin-Han Shiu

**Author notes:** Corresponding Author: Shin-Han Shiu, Michigan State University, Plant Biology Laboratories, 612 Wilson Road, Room 166, East Lansing, MI 48824-1312. Christina B. Azodi, Bioinformatics and Cellular Genomics, St. Vincent’s Institute of Medical Research, Fitzroy, Victoria, 3065, Australia. Contributions: CBA and SHS conceptualized the study. Data curation, formal analysis, implementation of the deep learning models, visualizations, and drafting the initial manuscript was done by CBA. JPL contributed to data curation and writing the machine learning pipeline. All authors contributed to writing the manuscript. All authors read and approved the final manuscript.

## Abstract

Plants respond to their environment by dynamically modulating gene expression. A powerful approach for understanding how these responses are regulated is to integrate information about *cis-*regulatory elements (CREs) into models called *cis-*regulatory codes. Transcriptional response to combined stress is typically not the sum of the responses to the individual stresses. However, *cis-*regulatory codes underlying combined stress response have not been established. Here we modeled transcriptional response to single and combined heat and drought stress in *Arabidopsis thaliana*. We grouped genes by their pattern of response (independent, antagonistic, synergistic) and trained machine learning models to predict their response using putative CREs (pCREs) as features (median F-measure = 0.64). We then developed a deep learning approach to integrate additional omics information (sequence conservation, chromatin accessibility, histone modification) into our models, improving performance by 6.2%. While pCREs important for predicting independent and antagonistic responses tended to resemble binding motifs of transcription factors associated with heat and/or drought stress, important synergistic pCREs resembled binding motifs of transcription factors not known to be associated with stress. These findings demonstrate how *in silico* approaches can improve our understanding of the complex codes regulating response to combined stress and help us identify prime targets for future characterization.

## INTRODUCTION

In order to survive and thrive, plants dynamically respond to changes in their environment. Given projected increases in global temperatures (1) and the frequency and severity of droughts, heat waves, and flooding (2, 3), improving our understanding of how plants regulate these dynamic changes will be critical for future efforts to breed and engineer more resilient crops (4) and for our ability to understand how a changing climate will impact diverse plant species (5). Most efforts to study stress response in plants have focused on how plants respond to a single stress in otherwise controlled conditions. However, in nature multiple stressors are typically present (6) and the response to combined stress may be different than the response to either of the stresses individually (7–9). For example, at the transcriptional level, ∼60% of *Arabidopsis thaliana* genes were found to respond to combined stress conditions in ways that are not predictable based on their responses to individual stressors (10). While efforts have been made to identify transcriptomic (7, 11, 12), metabolomic (13, 14), or physiological (9) changes in response to combined stress, the molecular mechanisms underlying how these complex changes are regulated remain unknown.

One major component regulating transcriptional changes to stress is the binding of one or more transcription factors (TFs) nearby a gene, which can change when and to what degree that gene is expressed. The importance of TFs for regulating transcriptional response to stress has made them targets for breeding and engineering plants for improved response to stresses, including salt (15), drought (16, 17), drought and heat (18, 19) stress. Further, genes underlying the domestication of crop species include TFs (20, 21). One approach to find the TFs driving stress induced changes in gene expression is to find the TFs associated with the non-coding regions of DNA near the transcriptional start site of a gene where TFs bind, or *cis-*regulatory elements (CREs). For hundreds of TFs in model species like *A. thaliana*, the DNA sequences that a TF can bind to (TF binding motifs; TFBMs) have been established *in vitro* (22, 23). In addition, putative CREs (pCREs) can be found computationally based on enrichment of specific *k*-mer sequences among co-expressed genes (24, 25). Previous studies have demonstrated that both known TFBMs and pCREs can be used to generate machine learning models that are predictive of a gene’s response to different environmental conditions (24, 26, 27). These predictive models are referred to as *cis-*regulatory codes. Nonetheless, current plant stress response *cis-*regulatory codes were built without considering additional factors that can also influence TF binding and transcriptional stress responses (28), including chromatin accessibility (29–32) and histone modifications (33, 34). Therefore, methods to integrate these additional types of omic information into the *cis*-regulatory codes are needed. In addition, at the physiological level, the effects of combined drought and heat are generally additive (35). However, it is unclear to what degree these responses are additive, synergistic, or antagonistic at the level of transcriptional regulation. Thus, *cis-*regulatory codes of these three different types of response patterns will be highly informative for understanding how they are regulated differently.

Here we explore the *cis*-regulatory codes of transcriptional response to single and combined heat and drought stress in *A. thaliana*. Heat and drought were selected because these stresses often co-occur in nature and elicit both overlapping and conflicting physiological responses in plants (36). Moreover, TFs and TF binding motifs are known for these stresses individually (37, 38). To better understand the regulatory logic underlying single and combined stress, first, we grouped genes likely to be co-regulated based on their shared pattern of transcriptional response under single and combined heat and drought stress (14) (Step 1, **Fig. 1**). Then, we used known TFBMs and enrichment based pCREs (Step 2, **Fig. 1**) to generate models of the *cis-*regulatory codes controlling these different patterns of responses to single and combined heat and drought stress using machine learning. To improve our models of the *cis-*regulatory codes and therefore our understanding of how response to single and combined stress is regulated in *A. thaliana*, we modeled regulatory interactions (Step 3A, **Fig. 1**), used a deep learning approach to integrate additional omics information (i.e. chromatin accessibility, sequence conservation, and histone marks) into our models (Step 3B,**Fig. 1**), and expanded the scope of our models by including pCREs identified outside of the promoter region (Step 3C, **Fig. 1**). In addition to providing a comprehensive overview of the *cis-*regulatory codes of response to single and combined heat and drought stress in *A. thaliana*, this study also exemplifies how a data-driven approach can be used to make novel discoveries in a complex system like gene regulation (Step 4, **Fig**.**1**).

**Fig. 1.**
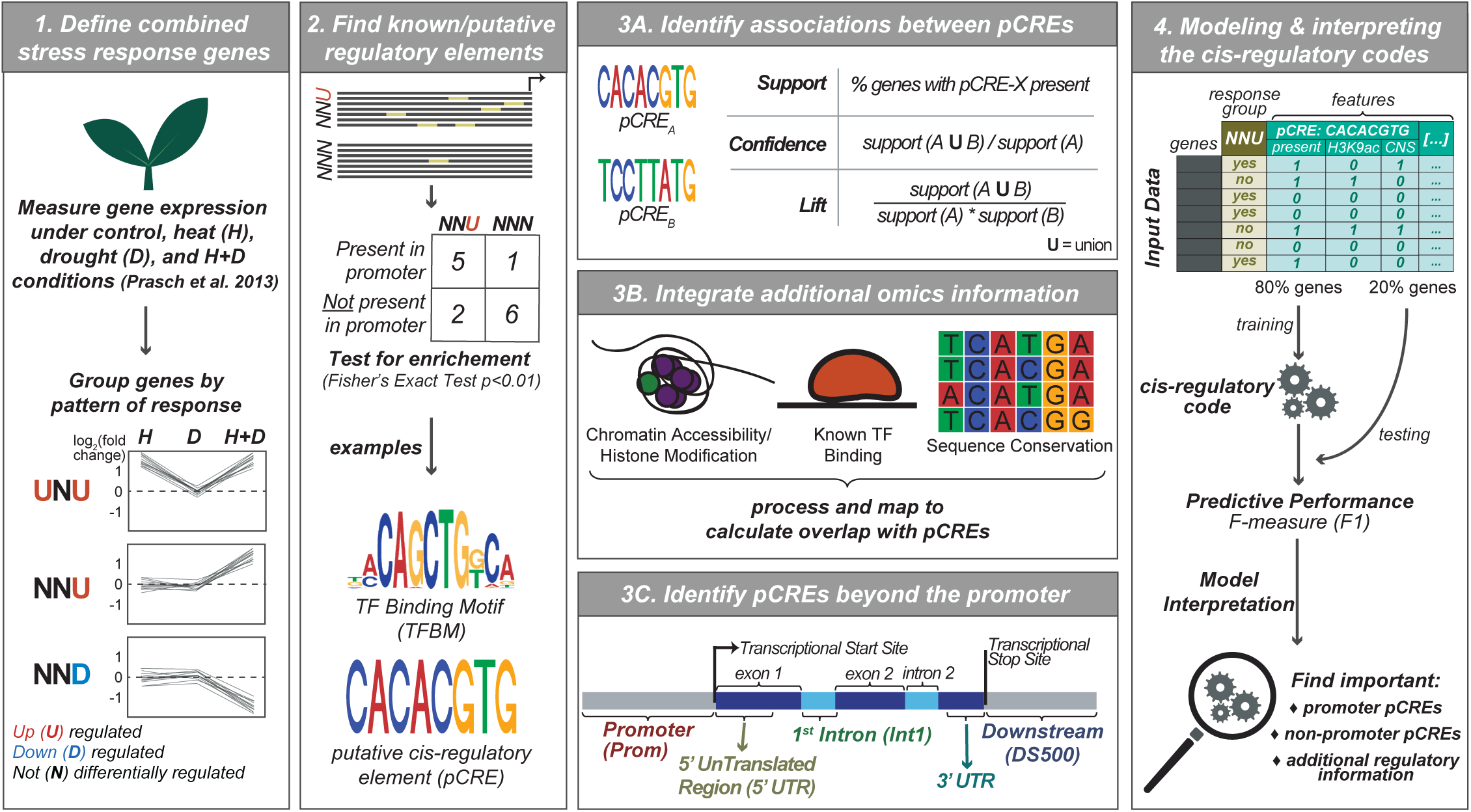
A framework for generating multi-omics models of the cis-regulatory codes. Step 1: Genes were grouped based on their pattern of differential expression under heat (H), drought (D), and H+D stress compared to control conditions. Step 2: For each response group, known TFBMs and putative cis-regulatory elements (pCREs) were identified based on site enrichment among response group genes (Fisher’s Exact Test; p-value < 0.01). Step 3: Information was gathered about associations between pCREs, their overlap with additional omics information, and pCREs located outside of the promoter regions. Step 4: All of this information was combined into machine learning models of the cis-regulatory codes and the models were interpreted to identify the most important components driving the predictions.

## MATERIALS AND METHODS

### Expression data processing, response group classification, and functional category enrichment analysis

Expression data for response to mild heat (32°C day/28°C night for 3 days), mild drought (30% field capacity), and combined heat and drought stress in *A. thaliana* were downloaded from NCBI Gene Expression Omnibus (GEO) (GSE46760) as normalized signal intensity values (14). The expression data was generated using the Agilent platform and probe data was converted into TAIR10 gene identifiers using IDswop from the “agilp” package in the R environment (39). If multiple probes were present for the same gene the mean of the probe intensities was used, unless the intensities were >20% different, in which case the gene was excluded. Differential expression folds and associated false discovery rate (FDR) adjusted *p-*values (i.e. *q-*values) (40) between each stress conditions and the control condition were calculated using limma (41) in the R environment.

Genes were classified as significantly up-regulated (U) if their log2 fold-change ≥ 1.0 with *q* ≤ 0.05, down-regulated (D) if their log2 fold-change ≤ −1.0 with *q* ≤ 0.05, or non-responsive (N) otherwise. Genes were clustered into “response groups” using a convention established by (10). Briefly, each gene was defined by its pattern of U, D, or N under heat, drought, and combined stress conditions. For example, a gene that is U under heat, D under drought, and N under combined stress was classified as in the UDN response group. To more clearly distinguish between genes belonging to a response group from genes that were considered non-stress responsive (NNN), genes were only considered NNN if they were not significantly differentially expressed (up- or down-regulated) with a log2 fold change cutoff of 0.8 under any of the three stress conditions or under any stress condition at any time point in the AtGenExpress database (http://www.weigelworld.org/resources/microarray/AtGenExpress/). P

Sequence data for the promoter, 5’ UTR, 3’ UTR, first intron, and downstream region for *A. thaliana* genes were downloaded from TAIR10. Genes whose promoter regions (1-kb upstream the transcriptional start site) overlapped with neighboring genes were excluded from the analysis. We tested if genes oriented in the same direction as their upstream neighboring gene were more likely to be correctly predicted than genes with partially overlapping promoter regions, but the results were not significant for most response groups (**Table S1**), so genes oriented in any direction were kept. For the analysis of the regulatory information in regions outside the proximal promoter, only genes that had sequence data available for all regions were included (**Table S2**)

The enrichment of GO terms (http://www.geneontology.org/ontology/subsets/goslim_plant.obo) and metabolic pathways (http://www.plantcyc.org) in the response group genes compared to NNN genes, were determined using the Fisher’s Exact test with *p-*values adjusted for multiple testing (42). As no AraCyc terms were enriched, only GO terms were discussed.

### Identification of known binding sites from in vitro TF binding data

Two sets of *in vitro* TF binding motif (TFBM) data were used to identify known binding sites. First, *in vitro* 200 bp binding regions for 344 TFs were collected from the DAP-Seq database (23). These 200 bp regions were derived from mapped sequencing peaks, and only peaks with a fraction of reads in peaks (FRiP) ≥ 5% were included. Second, position frequency matrices (PFMs) were obtained from the CIS-BP database for an additional 190 TFs without DAP-Seq data (22). CIS-BP PFMs were covered to Position Weight Matrices (PWM) adjusted for *A. thaliana’s* AT (0.33) and GC (0.17) background using the TAMO package (43). These 190 PWMs were then mapped to the putative promoter region (within 1kb upstream of the transcription start site) of *A. thaliana* genes using Motility with a threshold of *p*<1e-06 (http://cartwheel.caltech.edu/motility/). A gene was considered to be regulated by a TF if its putative promoter region overlapped with one or more known TFBM sites. We also identified a subset of known TFBMs that were enriched in the promoter regions of genes in a response group compared to non-responsive (NNN) genes using the Fisher’s Exact test (*p*<0.05), these TFBMs are referred to as the known enriched TFBMs (eTFBMs). To confirm that selecting eTFBMs was not resulting in overfitting, we repeated eTFBM finding for the smallest (NUN) and largest (UNU) response groups with 20% of the genes held out during the enrichment test and during model training (see below). This was repeated 100 times for each response group.

### Computational identification of novel pCREs and comparison with known TFBMs

To identify pCREs that were not covered by the available *in vitro* TF binding data, an enrichment based computational approach was taken (referred to as the iterative *k*-mer finding approach). With this approach, modified from (27), all possible *6*-mers tested for enrichment in the response group gene promoters compared to NNN gene promoters using the Fisher’s Exact test (*p*< 0.01). Multiple test correction was not used to avoid eliminating pCREs that may be important for a subset of genes in the response group. For *6*-mers that were enriched, their sequence was lengthened to all eight possible *7*-mers (e.g. ATATCG → <u>A</u>ATATCG, <u>T</u>ATATCG, <u>G</u>ATATCG, <u>C</u>ATATCG, ATATCG<u>A</u>, ATATCG<u>T</u>, ATATCG<u>G</u>, ATATCG<u>C</u>), which were then each tested for enrichment. The *k*-mer lengthening process continued until the longer *k*-mers were no longer significantly enriched. To confirm that the iterative *k-*mer finding approach was not resulting in overfitting, we repeated *k-*mer finding for the smallest (NUN) and largest (UNU) response groups with 20% of the genes held out during the enrichment test and during model training (see below). This was repeated 100 times for each response group. We also used the *k-*mer finding approach to find enriched pCREs in the 5’ UTR, 1^st^ intron, 3’ UTR, and 500 bp downstream region.

To assess the sequence similarity between (A) the pCREs identified for different response groups, (B) between the pCREs identified in different regions, and (C) between the pCREs and all known *in vitro* TFBMs, the Pearson’s Correlation Coefficients (PCC) between pCREs/TFBMs were calculated as in (26). PCCs between the top matching pCREs or pCREs/TFBMs are reported. Because the sequence similarity between top matching pCREs from (A) or from (B) would be greater by random chance if there were more pCREs in the comparison, the PCCs for (A) and (B) were reported as a percentile of a background distribution generated for each comparison based on a distribution of PCCs between top matching *6*-mers from groups of random *6-*mers of the same size as the groups in the comparison, repeated 1000 times. With this approach, the 50^th^ percentile indicates the similarity between pCREs from two response groups is no greater than random expectation, while a 99^th^ percentile would indicate the PCC is greater than 99% of the PCCs between random *6*-mers. To determine the degree of sequence similarity in (C), three PCC thresholds for each TFBM were calculated that range from least to most stringent. The lowest level of stringency is “better than random”, where the pCRE-TFBM PCC is ≥ 95th percentile of PCCs between the TFBM and 1,000 random *k*-mers. The next level of stringency is “between family”, where the pCRE-TFBM PCC is ≥ 95th percentile of PCCs between the TFBM and TFBMs from other TF families. Finally, the highest level of stringency is “within family”, where the pCRE-TFBM PCC is ≥ 95th percentile of PCCs between TFBMs from within the same family.

### Sequence conservation, chromatin accessibility, and histone mark data processing and analysis

Sequence conservation the between species conservation criteria, *A. thaliana* genomic regions that overlapped with ∼90,000 Conserved Noncoding Sequences (CNS) among 9 Brassicaceae species were used (44). DNase I Hyper-Sensitivity (DHS) regions were downloaded from GEO (GSE53322 and GSE53324) as peaks in bed format. These regions were identified from multiple tissues and developmental stages, including roots, root hair cells, leaf, seed coat, and dark grown *A. thaliana* Col-0 seedlings at 7-days old (45). Regions associated with activation-associated histone marks (H3K4me1: SRR2001269, H3K4me3: SRR1964977, H3K9ac: SRR1964985, and H3K23ac: SRR1005405) and with repression-associated histone marks (H3K9me1: SRR1005422, H3K9me2: SRR493052, H3K27me3: SRR3087685, and H3T3p: SRR2001289) were as compiled previously (46) using data from (47).

The percentage of times the sites of a pCRE overlapped with the 11 additional omics information (DAP-Seq, CNS, DHS, and eight histone marks) was calculated for each combination of pCRE and additional omics information for each response group. To determine how these overlaps were significant or not, 1,000 random, unique *6-*mers were generated and mapped to the promoter regions of response group genes, then the percentage of overlap with each combination of random *6-*mer and additional omics information was calculated for each response group. These overlap percentages were used to generate background distributions for overlap with each additional omic region, allowing us to convert the percent overlap scores for pCREs into percentiles along this background distribution. The percentage overlap with each additional omics information was also calculated for all CIS-BP motifs. Analysis of Variance (ANOVA), implemented in R v3.5.3, was used to determine if there were difference in the overlap percentage for each of the 11 additional omics information for each set of response group genes all pCRE, the top 10 most important pCREs (details below), the CIS-BP motifs, and the 1,000 random *6*-mers. The ANOVA *p-*values were adjusted for multiple testing (42). Finally, post-hoc Tukey tests, implemented using the HSD.test function from the agricolae package in R, were performed on comparisons with a significant ANOVA (*q*-value < 0.05) to identify which groups (i.e. pCREs, top 10 pCREs, CIS-BP, or random *6-*mers) had significantly different distributions in their percent of overlap with the additional omics information (*p <* 0.05).

To convert the additional omics information into features that could be used as input to our machine learning models, a new feature was generated for each pCRE – additional omics information pair (e.g. pCRE-DHS), where the value of the feature was set to 1 if the pCRE was both present in the promoter region of the gene and overlapped with the additional omics information and set to 0 if either or both of those criteria were not met. This resulted in a total of 12 features associated with each pCRE (i.e. the original presence/absence feature + the 11 additional features).

### Classic machine learning-based models of the cis-regulatory code

A classic machine learning algorithm called Random Forest (RF) (48) was used to generate models of the *cis-*regulatory code for each response group. These models were trained using a supervised learning approach, meaning they learned to predict the desired output (e.g. does the gene belong to response group NNU or NNN?) using example instances (i.e. genes) for which they have both the input features (e.g. presence of absence of pCRE-X) and the true classification (e.g. NNU or NNN). Different sets of input features were used throughout the study, including known TFBMs, promoter pCREs, combinatorial pCRE rules (see **Supplemental Methods**), overlap with additional omics information, and non-promoter pCREs.

RF was implemented using Scikit-Learn in Python 3 (49). To avoid training models that classify all genes as belonging to the more common response group, we balanced our input data by randomly down-sampling genes from the larger response group to match the number of genes in the smaller response group. Because the genes included in the input data can impact model training and performance, this process was replicated 100 times. To measure the performance of our models on a set of genes not seen by the model during training we used a 10-fold cross-validation scheme, where the input data was randomly divided into 10 bins, then a model was trained on bins 1-9 (i.e. the training set) and that model’s performance was measured based on how will it performed on the instances in the 10^th^ bin (i.e. the validation set). This was repeated, until each bin was used as the validation set one time. To select what values to use for two important RF parameters—maximum depth [3, 5, 10, 50] and maximum features [10%, 25%, 50%, 75%, 100%, square root(100%), and log2(100%)]—a cross-validated grid search implemented using GridSearchCV from Scikit-Learn was performed on the first 10 of the 100 balanced datasets (**Table S3**). The maximum depth parameter controls how deep each decision tree can be trained, where trees that are too shallow may not be able to capture complex patterns and trees that are too deep may overfit, meaning they would predict the training genes well, but would not generalize to predict genes not included in training well (e.g. the validation set or new genes). The maximum features parameter controls how many of the input features each decision tree in the forest will be allowed to use, where too few will result in poor performance from individual decision trees and too many will result in most decision trees in the forest identifying the same pattern.

Model performance was evaluated using the F-measure (F1) (50), or the harmonic mean of precision (True Positive / True Positives + False Positives) and recall (True Positives / True Positives + False Negatives), where an F1=1 would indicate all gene were perfectly classified, and an F1=0.5 would indicate the model did no better than random guessing. Model performance was compared using two-sided paired t-tests, with response groups paired (n=7). For each model we also determined which genes were correctly classified as belonging to a response group, *R*. Every balanced run of the model could have predicted a different subset of genes as belonging to *R*. Thus, a final classification call that a gene, *G*, belongs to group *R* was determined if the mean predicted probability of 100 balanced runs ≥ the predicted score threshold (i.e. the threshold between 0 and 1 that maximized model performance averaged over replicates). For each balanced run, we identified the predicted score that maximized the F1. We took the average of the predicted score maximizing F1 for all 100 runs as the predicted score threshold. Then, models with similar F1 scores could be compared to see if they predicted a different subset of genes. Finally, the relative importance of each feature in a RF model was determined using the importance score function built into the Scikit-Learn implementation of RF. This function calculates feature importance as the normalized decrease in node impurity across the decision trees when that feature is used to divide a node, known as the Gini Importance (48). To confirm our eTFBM and pCRE features were not overfit, we trained RF models using the eTFBMs and pCREs identified with 20% of the genes held out as features using the genes not held out as our training instances. After the models were trained, they were applied back to the held-out 20% of genes and the performance (F1) was calculated on the held-out genes only.

### Convolutional neural network-based models of the cis-regulatory code

Convolutional neural networks (CNNs), a deep learning algorithm (48), were tested to see if it could better integrate additional omics information into our models of the *cis-*regulatory code. CNNs were implemented in Python 3.6 using Tensorflow 2.0 (51). CNN models were made up of four layers: input, convolutional, dense (i.e. fully connected), and the output (i.e. the prediction). The input is a 3-dimensional array [rows x columns x layers] where each layer contains data from a different gene, each column (size= # of pCREs for that response group) contains different pCREs, and each row (size=12) contains either pCRE presence/absence or overlap with additional omics information. The convolutional layer is composed of kernels (i.e. pattern finders) with the dimensions [12 x 1], using a stride length =1, this resulted in each kernel passing over each pCRE one time and resulting in an output with dimensions [# kernels x # pCREs]. The starting kernel weights were initialized randomly and were scaled relative to the size of the input data using Xavier Initialization (52). The output from the convolutional layer was flattened (i.e. changed the output from a 2D array to a 1D array with shape [1 x (# kernels x # pCREs)]) and then passed to the dense layer. A non-linear activation function (rectified linear units; ReLU) was applied to both the convolutional and dense layers, and a sigmoid activation function was applied to the final output layer to facilitate making a binary decision (e.g. NNU vs. NNN). Weights were optimized using the Stochastic Gradient Descent with momentum (SGDm) (momentum=0.9) as implemented in Tensorflow.

Three strategies were used to reduce the likelihood of the CNN models overfitting, where models train so specifically to the training data that they do not generalize well to new data. First, L2 regularization was applied to the kernel weights in our convolutional layer, forcing the weights to shrink toward zero. Second, dropout regularization was applied to the dense layer, meaning during each iteration of training a random subset of the dense nodes were removed. This essentially adds randomness to the model and encourages the network to learn more general patterns in the data, rather than specific ones that may be overfit. Finally, CNNs can overfit to the training data if they are allowed to train for too many iterations. However, training for too few iterations will result in a model that has not yet converged (i.e. underfitting). To determine when to best stop training, we used an early stopping approach implemented in Keras (https://keras.io/callbacks/#earlystopping), where the training data was further split into training (90%) and validation (10%) and training stopped when model performance had not increased (min_delta = 0) for 10 iterations (patience = 10) on the validation data, with the maximum number of training iterations limited to 1,000. As with the RF models described above, CNN models were trained on balanced datasets. Because of the greater computational power needed by CNNs, instead of the cross-validation approach used for RF, the balanced data was divided into a training set (90%) and testing set (10%) and performance was measured on the testing set. This was repeated 100 times using different training and testing sets for each replicate. Model parameters were selected using a random search across the parameter space with five-fold cross validation with ∼4,800 iterations (implemented using RandomizedSearchCV in Scikit-Learn). Parameters in the search included the learning rate, the number of kernels in the convolutional layer, the number of nodes in the dense layer, the dropout rate, and the L2 regularization rate (see **Table 7**).

The importance of each pCRE and its associated additional omics information was determined by measuring the difference in model performance between the original model and a new model when the values in all rows for a pCRE column were set to zero (i.e. not present and not overlapping with the additional omics information) for all genes. Thus, larger positive differences indicate pCREs were important. Negative scores indicate zeroing out the pCREs in question actually improved model performance. The change in performance measured using the area under the receiver operator characteristic, rather than the F1 because it does not require the selection of a classification threshold. The median importance scores across the 100 replicates were used to summarize the importance of each pCRE and its associated additional omics information. To determine what patterns the CNNs learned to identify, we extracted the weights from each kernel in the convolutional layer of our trained CNN models. Given that each of the 100 replicates involved training a CNN model with either 8 or 16 kernels (see **Table 7**) we had had between 800-1,600 trained kernels for each model of the *cis-*regulatory code. To summarize this information we used hierarchical clustering with dynamic branch cutting (minimum cluster size = 250) to group kernels based on the similarity of their weights and found the median weight at each position for each cluster. Kernel importance was measured as described above, where the change in model performance after a kernel’s weights were set to zero (i.e. identifying no pattern) was calculated for each kernel. The median kernel importance scores across all kernels in a cluster are show.

### Availability of data and materials

The datasets supporting the conclusions of this article are available as described by the original authors (14, 22, 23, 44, 45, 47). All code needed to reproduce the results from this study are available on GitHub (https://github.com/ShiuLab/Manuscript_Code/2019_CRC_HeatDrought). This repository also contains a detailed README.md file which describes our analyses in more detail, provides the commands used to generate the results in this study, lists additional software needed, and includes links to the most recent versions of the scripts used. Scripts are implemented in Python or R.

## RESULTS

### More than 50% of stress responsive genes have unpredictable responses to combined heat and drought stress based on single stress response

In order to study the regulation of transcriptional response to single and combined stress, we first identified groups of genes that were likely to be co-regulated based on their shared pattern of transcriptional response to three stress conditions: heat, drought, and combined heat and drought stress using transcriptome data from an earlier study (14). Transcriptional response was indicated with one of three abbreviations based on up-regulation (U), down-regulation (D), or no response (N), and response categories were labeled with a three-letter designation, where the first, second, and third letter indicated response to heat, drought, and combined stress, respectively. For example, genes that were up-regulated under heat and combined stress, but not under drought alone were placed in the UNU response group. These response groups were further categorized based on if the response to the combined stress was similar to (“independent”: UNU, NUU, DND, or NDD), less than (“antagonistic”: UNN, NUN, DNN, or NDN), or greater than (“synergistic”: NNU or NND) the sum of the responses to the single stress conditions (**Fig. 2A**).

**Fig. 2.**
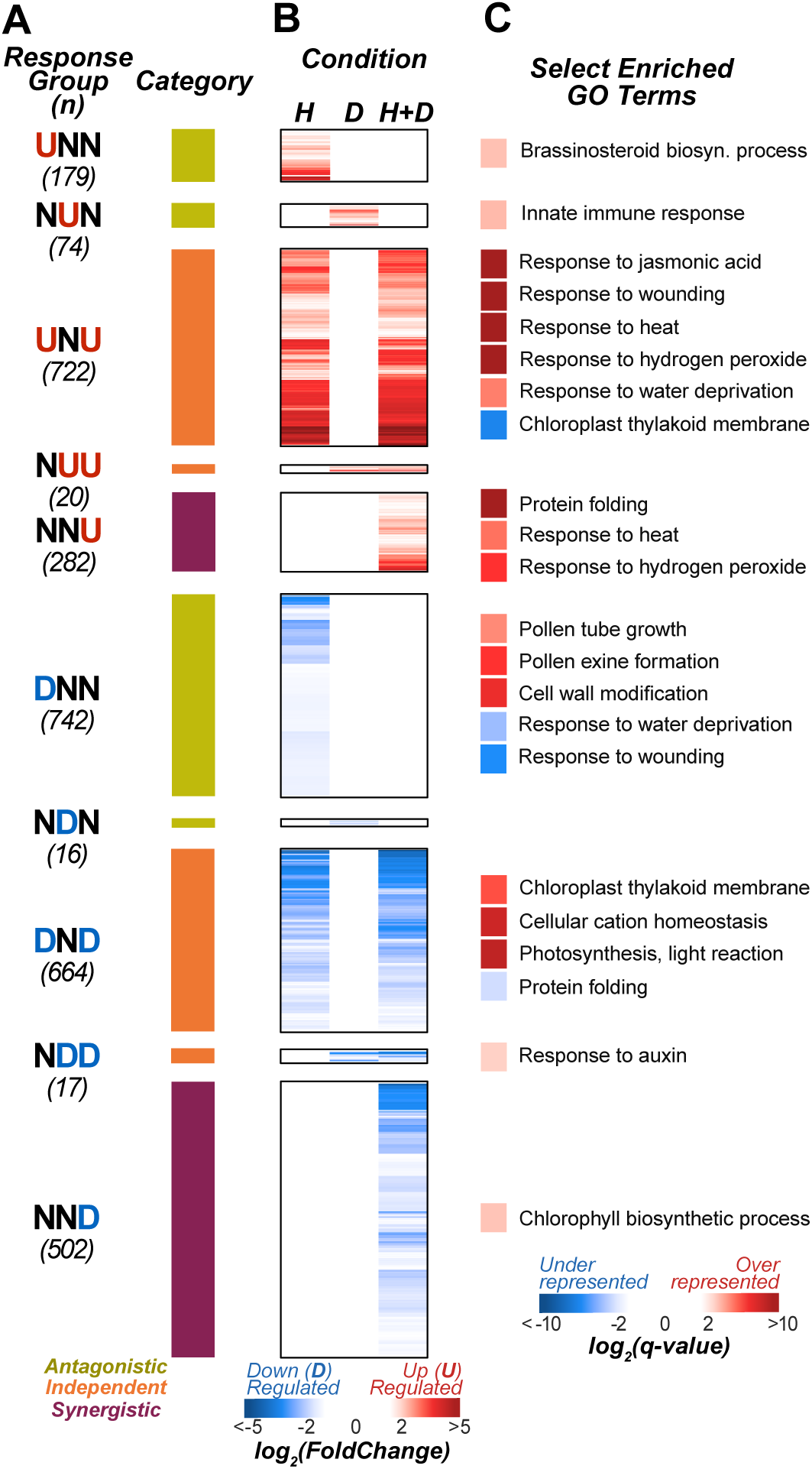
Gene expression response groups for single and combined heat and drought stress. (A) Gene expression response groups included in the study where the three-letter response codes signify up-regulation (U), down-regulation (D), and no significant change in expression (N) ordered based on response to heat, drought, and both stresses. The number below the response group name is the number of genes in that response group that have non-overlapping promoters (1kb upstream of TSS) with neighboring genes. Colored bars designate if genes in the response group are considered to have antagonistic (yellow), independent (orange), or synergistic (purple) responses to combined stress. (B) The log2 Fold Change in expression under heat (H), drought (D), and H+D compared to control for each gene (X-axis), sorted by response group. If the absolute value of the Log2(FC) ≤ 1, colored white (N). (C) Select Gene Ontology (GO) categories that were enriched for genes belonging to the different response group compared to all other genes. GO categories with a large positive log2(q-value) (red) are over-represented, while those with large negative log2(q-value) (blue) are under-represented in that response group.

Among genes that were responsive to at least one stress (n=3,218), 43%, 29%, and 24% genes were in the independent, antagonistic, and synergistic response groups, respectively (**Fig. 2B; Table S1**). The remaining 4% of genes belonged to rare response groups (e.g. DUN, UUD) and were not considered in our analysis. Most of the genes in the independent and antagonistic response categories were responsive (up- or down-regulated) to heat, rather than drought stress. The dominance of the heat response could be due to: (1) the mild nature of the experimental drought stress (14), (2) a possible overriding influence of heat stress, e.g. heat response dominates over salt stress (10), or (3) the fact that the expression data is derived from leaf where drought-induced osmotic stress has a lesser effect compared to root (53). Gene Ontology enrichment analysis (**see Methods**) confirmed that different response groups are enriched for genes with different biological functions (**Fig. 2C; Table S4; Supplemental Info**). Further, this analysis demonstrated that genes in the synergistic response groups tended to overlap functionally with genes in independent response groups. For example, both up- regulation independent (UNU) and synergistic (NNU) response groups were enriched for response to heat and hydrogen peroxide. This reinforces the idea that genes with similar biological functions are not necessarily co-regulated.

In summary, we found that ∼55% of genes responsive to at least one stress showed either antagonistic or synergistic responses to combined heat and drought stress. Genes in these non-additive response groups are especially interesting because knowing how they respond to heat stress and drought stress independently does not help us predict how they will respond to combined stress. Because these non-additive responses to combined stress were so prevalent, we hypothesized that unique regulatory codes must exist that are able to fine tune transcriptional response under combined heat and drought stress.

### Combinatorial stress response patterns can be predicted using known and putative regulatory elements

Because TFs and associated binding sites regulating combinatorial stress response are unknown, we set out to identify responsible TFs by taking advantage of available *in vitro* TF binding region and motif (known TFBMs) data for 344 TFs from the DAP-seq (23) and CIS-BP (22) databases. First, 197 of the 344 known TFBMs were identified as enriched in the promoter region of at least one set of response group genes (*p*<0.05; referred to as enriched TFBMs, eTFBMs, see **Methods**). On average, response groups were enriched for 35 known TFBMs (range: 0-87) from 27 TF families (referred to as enriched families, **Table S1**). In parallel, to identify regulatory sequences not covered by known TFBMs, we searched for putative *cis-*regulatory elements (pCREs) by identifying *k*-mers enriched in the promoter regions of genes in each response group compared to genes not responsive to stress (see **Methods**). Response groups were enriched for 68 pCREs on average (range: 7-158).

To determine the extent to which known eTFBMs and co-expression-based pCREs can explain combined stress response patterns, we used the presence or absence of eTFBM and pCRE sites as features (i.e. independent variables) in Random Forest (RF) models to classify genes as belonging to a response group or as non-responsive under any stress condition (i.e. the dependent variable). Because machine learning models need to learn from sufficient training data, only response groups with >20 genes were used. Model performance was measured by calculating the F-measure (F1) on a set of data held out from model training, where an F1=1 would be a perfect classification and an F1=0.5 would be no better than random guessing (see **Methods**). Both the eTFBM and pCRE-based models were able to predict single and combined stress response groups better than random guessing (**Fig. 3A**). However, models built using pCREs (median F1_pCRE_=0.64) significantly outperformed those built using known eTFBMs (median F1_eTFBM_=0.58; paired t-test, *p*=3.7×10^−4^). One concern was that our models may be overfitted because pCREs and eTFBMs finding was performed using all genes in a response group (e.g. all NNU and NNN genes). To test this, we repeated the pCRE and eTFBM finding and RF model training/cross-validating on 80% of the genes and then applied and measured the performance of the models on the remaining 20% of genes. This was repeated 100 times for both the largest (UNU) and smallest (NUN) response groups and no significant difference in performance was detected (paired t-test, *p*=0.22-0.99; **Table S5**), indicating our models were not overfitted. Further, when we used all known TFBMs (i.e. both enriched and non-enriched), the model performance decreased further (median F1_TFBM_=0.54). These findings support the notion that pCREs contain additional omics information not captured by the TFBM data. This is not to say that pCREs can completely replace TFBM data because models built using the enriched TFBMs and pCREs were able to correctly classify different subsets of genes (**Fig. S1**). However, including both types of elements as features did not improve model performance compared to only using pCREs (median F1_pCRE+eTFBM_=0.64; paired t-test, *p*=0.51). Thus, we choose to focus on pCRE based models for the remainder of the study.

**Fig. 3.**
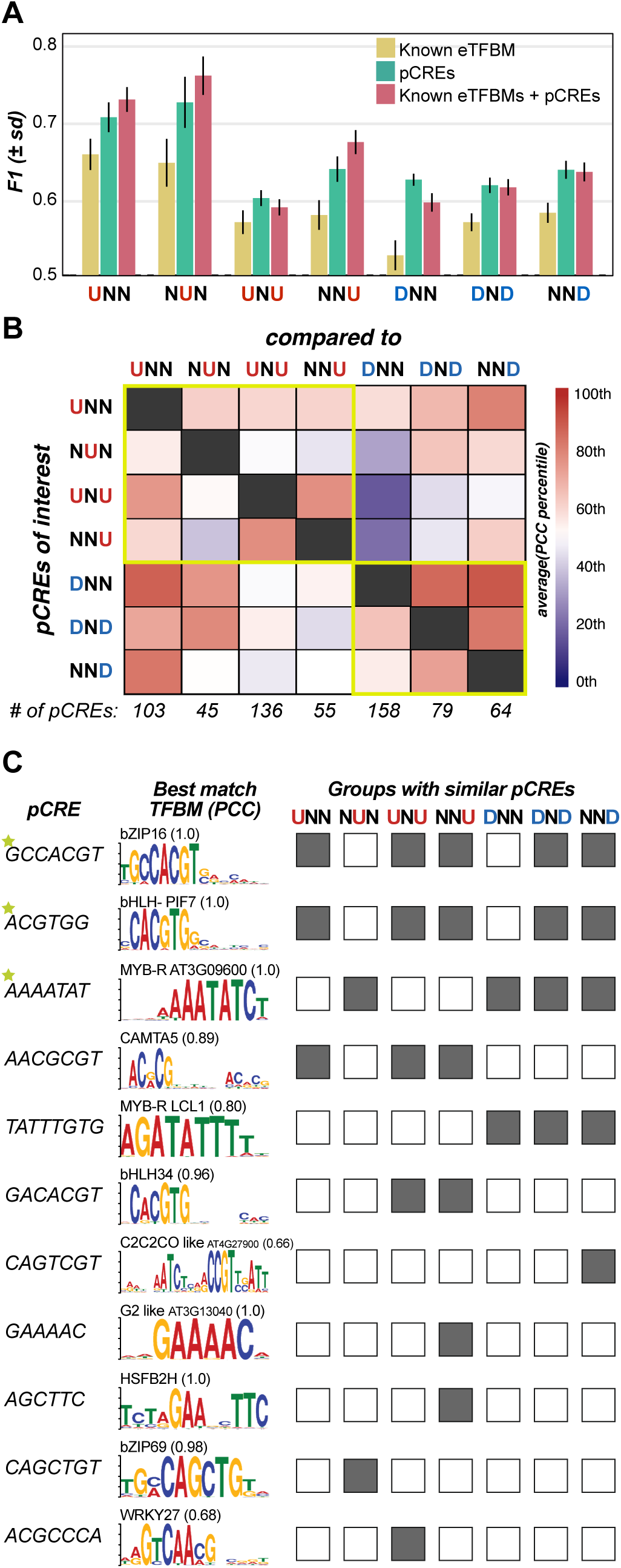
Known TFBM and pCRE models of the cis-regulatory codes. (A) Predictive performance (F1) of Random Forest machine learning models using known TFBMs (yellow), pCREs (teal), or both (rose) as input features as input for predicting response group vs. non-responsive genes. (B) Average sequence similarity (PCC) percentile between pCREs from the response group on the X-axis and their top matching pCRE from the response group on the Y-axis. The percentiles were calculated for each comparison based on a distribution of PCCs between top matching 6-mers between groups of the same size as the response groups in the comparison, where 50th percentile indicates the similarity between pCREs from two response groups is no greater than random expectation. (C) Select example pCREs (first column) and a motif logo of the known TFBM the pCRE is most similar to. The similarity scores (Pearson’s Correlation Coefficient; PCC) between each pCRE and its best match are shown. The pCREs are sorted from most to least commonly enriched across response groups, where the boxes indicate what response groups the pCRE was enriched (gray) or not enriched (white) in.

Next, we quantified the degree of sequence similarity between pCREs identified for different response groups to assess how the *cis-*regulatory programs differs across response groups. To account for different response groups having different numbers of pCREs, the PCC between the top matching pCREs from two response groups was reported as the percentile of a background distribution generated from the PCC between top matching 6-mers from groups of 6-mers of the same size as the number of pCREs for each response group. Using this approach, the average pCRE percentile overlap ranged from 24^th^ to 80^th^ between response groups (mean = 57^th^ percentile; **Fig. 3B**), with response groups that share the same direction of response (yellow boxes) being more similar to each other than response groups that respond in different directions (e.g. UNU vs DNN) (ANOVA; *p*<1×10^−4^). Interestingly, of the pCREs found among the most response groups, the top three, GCCACGT, ACGTGG, and AAAATAT (stars, **Fig. 3C**) were significantly similar to TFBMs associated with circadian clock TFs bZIP16, PIF7, and RVE8, respectively (54, 55). PIF7 has been shown to negatively regulate *DREB1* as a means to avoid hindering plant growth by the accumulation of DREB1 when the plant is not under stress (56). Our findings further support earlier studies that stress response regulation has a significant circadian clock component (57).

In summary, the *k*-mer finding approach identified pCREs that, when used as predictive features, were better able to classify genes by their response groups than known enriched TFBMs. Further, while some pCREs were identified across multiple response groups, the fact that average pCRE similarity between response groups was only the 57^th^ percentile, suggests there are substantial regulatory differences between the different responses to single and combined heat and drought stress. Finally, while we were able to classify genes by their response group well above random expectation (median F1_pCRE_=0.64), there was still ample room for model improvement. Because TFs frequently work in concert to regulate gene expression (58, 59), we first incorporated interactions between TFs into our models by identifying interactions between pCREs. We identified interactions between pCREs for each response group using two statistical approaches: association Rule and iterative RF. However, pCRE pairs identified did not improve model performance when used as features alone or with pCREs (**Fig. S2** and **Supplemental Info**), unlike in high salinity stress (26). Thus, we next explored improving our models by integrating additional types of omics information and including pCREs located outside the proximal promoter.

### Additional omics information can improve models of the cis-regulatory codes

To account for additional levels of regulation involved in response to single and combined heat and drought stress, we next explored adding chromatin accessibility, histone modification, sequence conservation, and known TF binding sites data to our models of the *cis-*regulatory codes. We included information about chromatin accessibility from DNase I Hypersensitive Sites (DHS) (27, 60) and eight histone marks (ChIP-seq) (47, 61, 62) because both can impact the TF binding. In addition, information about sequence conservation across the *Brassicaceae* family (Conserved Noncoding Sequences: CNS) was included as true CREs may be under selection and therefore may be more likely to be conserved (63, 64). Finally, *in vitro* TF binding regions identified in *A. thaliana* (described above) (23) were also included. These data are collectively referred to as “additional omics information”.

To determine if additional omics information would improve our understanding of the *cis*-regulatory codes of combined stress response patterns, we included these data as features (see **Methods**) in our RF models and assessed whether their inclusion improved predictive performance. While models utilizing this additional omics information improved the average performance for a few response groups (i.e. NNU, DNN), overall, they did not perform significantly better than pCRE-only models (median F1_pCRE+ARI_=0.66; median F1_pCRE_=0.64; paired t-test, *p=*0.062) (olive; **Fig. 4A**). One possible reason for this lack of improvement could be that RF, while robust at dealing with heterogeneous input data (e.g. multi-omics data), struggled to learn predictive patterns in our data because it treats each input feature as an independent piece of information, even when they are not. For example, each decision tree in a RF model only had access to a subset of the features for training, and that subset was selected randomly, without any consideration of associations between the features. Because our additional omics information features each provide more information about a particular pCRE, they are not independent. Further, assessing the whole profile of additional omics information associated with a pCRE could uncover new predictive patterns.

**Fig. 4.**
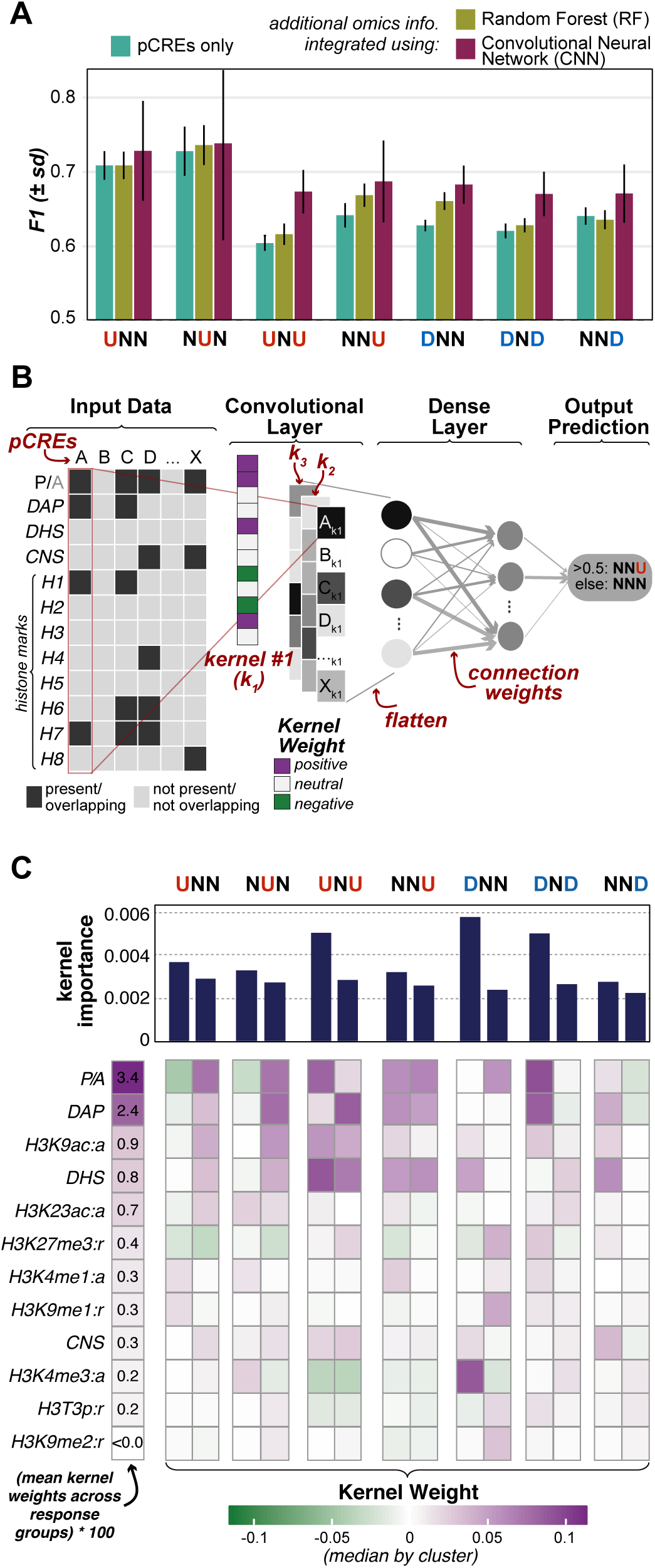
Integration of additional omics information into models of the cis-regulatory codes. (A) Predictive performance (F-measures (F1)) of Random Forest models using pCREs (teal, same as in Fig. 3A) and pCREs + additional omics information (olive) and of Convolutional Neural Network (CNN) models using both pCREs + additional omics information (rose). The larger error around NUN models is due to the small number of NUN genes available for model training. (B) An illustration of the internal workings of the CNN models and how the trained kernels (i.e. pattern identifiers) in those models were used to understand the patterns of additional omics information the models were trained to identify. (C) Summary of results from interpreting the trained CNN models. The feature types (i.e. presence/absence (P/A) and additional omics information) were sorted based on the average kernel weights across all kernels trained for all response groups and replicates (first column). The remaining columns represent kernel clusters for specific response groups. For each response group, all trained kernels from all CNN replicates were clustered using hierarchical clustering with dynamic cutting (min cluster size=250 kernels). The median kernel weights and kernel importance scores are shown here for the two clusters with the highest median kernel importance for each response group. Large kernel weights (dark purple) indicate the presence of that pCRE or its overlap with the additional omics information was an indicator of belonging to the response group rather than NNN. Note that some of the most important kernel clusters had negative weights for pCRE presence/absence. These kernels likely trained to learn patterns associated with the non-responsive gene group (i.e. NNN).

To address the limitation of RF, we applied a deep learning approach: convolutional neural networks (CNNs). CNNs are frequently used in image classification because when given training data (e.g. many photographs of cats) they are able to learn local patterns called kernels (e.g. triangles that resemble cat ears) and associate those kernels with what is being predicted (e.g. is there a cat in the photograph). However, they have been applied successfully to study genomic data (65, 66). We hypothesized we could train CNN models to look for patterns in the additional omics information available for each pCRE and to then associate those patterns with a response group (**Fig. 4B**; see **Methods**). Using this approach, our ability to predict response groups increased (median F1_CNN_=0.68) compared to the pCRE only models (median F1_pCRE_=0.64; paired t-test, *p*=0.002), a 6.2% improvement in the median F1, with the largest improvements for the UNU, DNN, DND, and NNU response groups (where F1 increased by ≥ 0.05) (rose; **Fig. 4A**).

### Interpreting deep learning models provides insight into the cis-regulatory code

To understand what combinations of additional omics information were important for the ability of our CNN models to classify genes by their response group, we interpreted our CNN models by visualizing the trained kernels and measuring their importance. During the process of model training, each kernel learns a particular “pattern”, i.e., how much value, or weight, should be given to each feature to best predict if a gene belongs to a response group. In the example shown in **Fig. 4B**, kernel #1 (k_1_) trained to look for pCREs that were present and that overlapped with a DAP site and with histone marks for H1 and H7 (positive kernel weights), but not H4 or H6 (negative kernel weights). Then, each trained kernel scans across the input data and generates an output value for each pCRE based on how well it matches the pattern. For example, when k_1_ was used to scan pCRE-A through pCRE-X, it led to a large (i.e. dark) value for pCREs that match its pattern (e.g. pCRE-A) and a small value (i.e. light) for pCREs that do not match its pattern (e.g. pCRE-D). To assess which types of features were most important (i.e. highest weighted) among kernels from CNN models for each response group, we extracted the trained kernels (i.e. a list of 12 weights) for each kernel in each replicate, clustered them into groups with similar patterns of weights, and calculated the median weight assigned to pCRE presence/absence and each additional omics information for each cluster (**Fig. 4C, S3**; see **Methods**).

To measure the overall importance of each kernel, we calculated the change in model performance on the test data (i.e. data not used for training) when each kernel was zeroed out (i.e. all weights set to zero; see **Methods**). We then reported the median kernel importance for each kernel cluster (**Fig. 4C, S3**). For example, when a kernel in the first kernel cluster for DNN was set to zero, model performance (measured using the area under the receiver operator characteristic; see **Methods**) dropped by > 0.005. Note that the performance decreases are all very small, indicating the models were robust to perturbation likely because more than one kernel was trained to learn important patterns. Overall, the presence or absence of the pCREs (P/A) had the highest median weights (leftmost column; **Fig. 4C**). Of the additional omics information, DAP, H3K9ac, and DHS had the next highest kernel weights, suggesting known TF binding, the acetylation of lysine 9 on histone H3 (a hallmark of active promoters (67)), and chromatin accessibility were consistently useful features for predicting response to single and combined stress. Other types of additional omics information were weighted differently in important kernel clusters for different response groups (second column and on; **Fig. 4C**). This was especially true of histone mark features. For example, H3K27me3 tended to be negatively weighted in important kernel clusters for up-regulation response groups (UNN, NUN, NNU) but neutral or positively weighted in important kernel clusters for down-regulated response groups (DNN, DND). Together with the fact that H3K27me3 is known to be associated with gene silencing (68), this finding suggests that lysine 27 trimethylation may play a role regulating response to single and combined heat and drought stress. However, we also found that H3K4me3 had a large positive weight for the most important DNN kernel cluster and negative weights for some up-regulation clusters (UNU, NNU). This was unexpected given that H3K4me3 is associated with active promoters (68).

In summary, we found that the integration of additional omics information into our models of the *cis-*regulatory codes using CNNs improved our ability to classify genes by their pattern of response to single and combined stress. While some information (e.g. TF binding, H3K9ac) was important for all response groups, other information (e.g. H3K4me3, H3K27me3) was only important for one or a few response groups, indicating that different response groups may be subject to distinct epigenetic regulatory signals. The usefulness of these data was especially surprising given some of the limitations of the data. For example, most of the data were generated either *in vitro* (e.g. DAP) or under growth conditions that do not match the transcriptome data used for this study (e.g. DHS, histone ChIP-seq).

### pCREs identified outside the promoter region are predictive of response patterns

The models discussed thus far were based on features located in the proximal promoter regions typically housing regulatory sequences in plants (69). However, plant regulatory sequences can also be located in the 5’ untranslated region (5’ UTR) (70), first intron (Int1) (71), and 3’ UTR (72). We also cannot rule out that some regulatory sequences can be present downstream of the transcriptional stop site (DS500). To assess the extent to which pCREs outside of the promoter regions were predictive of combined stress response patterns, the iterative *k*-mer finding approach was repeated in the 5’ UTR, Int1, 3’ UTR, and DS500. Then, predictive models were built using either pCREs from each region individually or in combination as features. Because sequence information was not available for all five regions for all genes (particularly 5’ and 3’ UTRs), we removed between 47 and 587 genes from each response group to make our models comparable. Importantly, this means that the performance results from our earlier machine learning models would not be directly comparable. In order to establish a direct comparison, we also re-ran the iterative *k*-mer finding and modeling on the promoter region using the smaller subsets of genes.

Models built using pCREs located in promoter or, surprisingly, DS500 regions outperformed models built with pCREs from other regions (*Tukey test*; **Fig 5A**). DS500 pCREs substantially outperformed promoter pCREs for the NUN response group in terms of F1 (+0.06, **Fig 5A**), as it correctly classified 2 more genes and reduced the false positives by 14 (**Fig S4**). Interestingly, the most predictive DS500 pCRE, ACTTTG, shares significant sequence similarity (PCC=0.92) with the known TFBM for WRKY46, which has known roles in drought response. This pCRE was not enriched in the promoter region, emphasizing the potential importance of the DS500 region for *cis* regulation. Although the 5’UTR and 3’UTR pCREs did not perform as well as those in promoters and DS500s, they were significantly better than random expectation (t-test: *p=*0.02, 0.006, respectively), however Int1 pCREs were not significantly different than random (p=0.75). Because models built using pCREs from different regions were able to correctly classify different subsets of genes (**Fig. S4**), we used pCREs from all regions as features and the resulting models (the ALL column, **Fig. 5A**) outperformed all single region-based models, suggesting that pCREs located beyond the promoter region are important for regulating combined stress response.

**Fig. 5.**
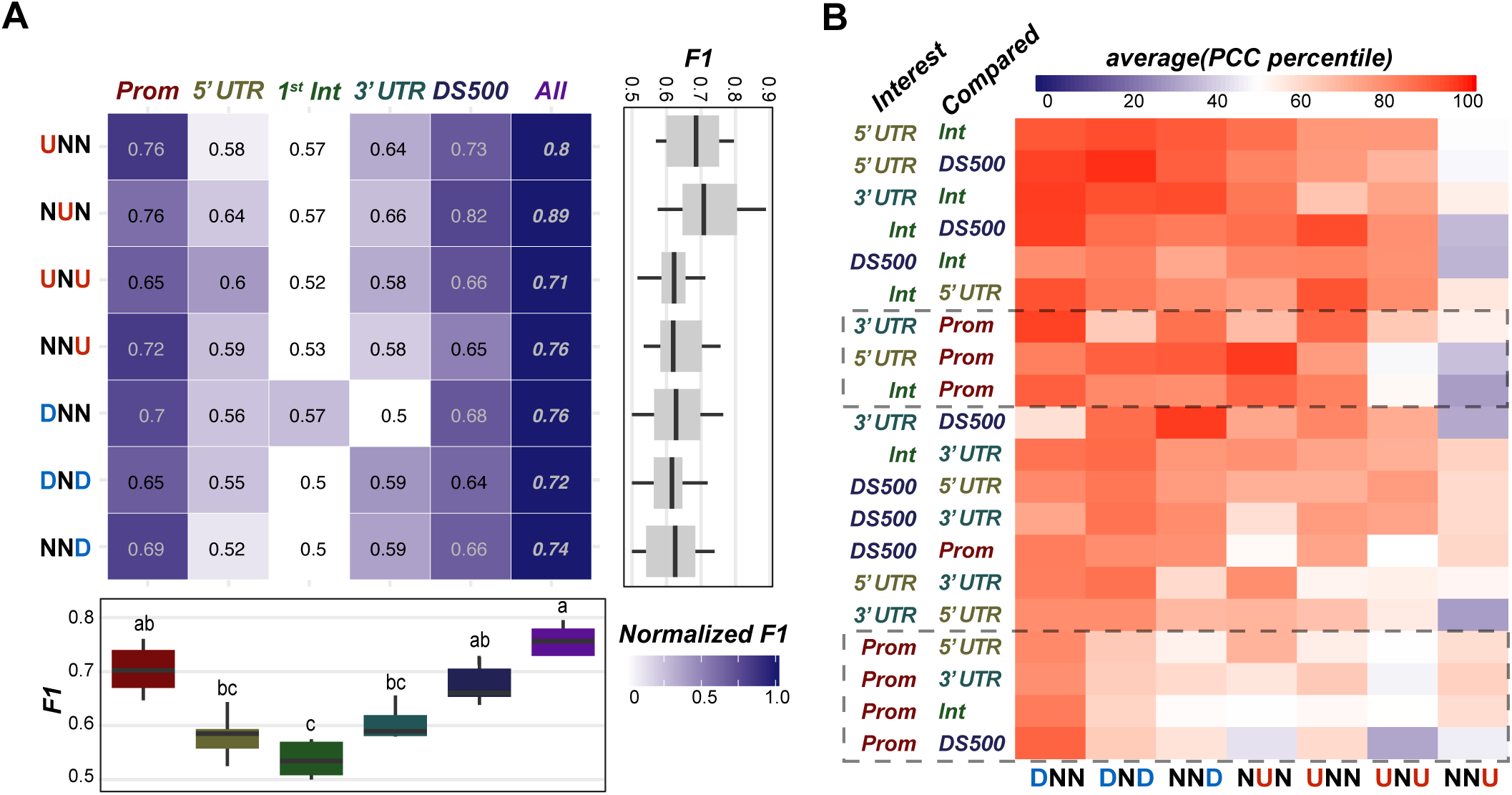
Promoter and non-promoter pCRE based models of the cis-regulatory codes. (A) Predictive performance (F1) from Random Forest models using pCREs found in the promoter, 5’ UTR, first intron (1st Int), 3’ UTR, downstream region (DS500), or all regions (All) as input features. The box color represents the F1 scores normalized by the F1s of each response group (the darkest blue represents the best set of input feature for each response group) with the actual F1 provided in each box. The boxplot shows the distribution of F1 scores for each region (below) and for each response group (right). Letters on the top of boxplots signify significant differences by region based on the Tukey test (p<0.05). (B) Average sequence similarity (PCC) percentile between pCREs identified from a region of interest (left column) compared to their top matching pCRE from another region (right column). The percentiles were calculated for each comparison based on a distribution of PCCs between top matching 6-mers between groups of the same size as the regions in the comparison, where 50th percentile indicates the similarity between pCREs from two regions is no greater than random expectation.

To determine if the pCREs identified from different genetic regions were unique to that region or found across regions, we calculated the similarity percentile between the best matching pCREs between regions within a response group as we did above with promoter pCREs from different response groups (see **Fig. 3B**). Overall, pCREs from different regions were more similar to each other than would be expected by random chance (>50^th^ percentile, red; **Fig. 5B**). This was especially true for down-regulation response groups, suggesting that regulatory elements involved in down regulating genes are either less region specific or are more likely to be located in multiple regions around the gene. Interestingly, the only response group where this was not the case was NNU, where the average pCRE similarity between regions was frequently near or below the 50^th^ percentile. Given the promoter pCREs were the most predictive of NNU, this suggests the regulatory circuitry for synergistic up-regulation is specific to the promoter region. Finally, we observed that while non-promoter pCREs tend to be similar to promoter pCREs (top horizontal box), the promoter pCREs were less similar to non-promoter pCREs (lower horizontal box). This indicated that promoter-specific pCREs are common, while pCREs identified in regions outside the promoter tend to be found more universally around the genes in a response group.

In summary, incorporating pCREs identified outside of the proximal promoter region improved our ability to predict response to single and combined heat and drought stress. Of the five regions assessed, the DS500 pCREs performed marginally better than promoter pCREs for two of the seven response groups. Taken together, this suggests that while most of the pertinent regulatory information is in the promoter regions, CREs important for response to single and combined heat and drought stress may be located outside the promoter region.

### Characterizing the most important pCREs identifies key features of the combined heat and drought stress cis-regulatory codes

We have demonstrated that adding multi-omics data and expanding our search for putative regulatory elements beyond the promoter region has improved our models of the *cis-*regulatory codes. While these models are still not perfect, they perform well above random expectation and therefore can be used to illuminate the *cis-*regulatory codes of response to single and combined heat and drought stress in *A. thaliana*. To this end, here we further characterize a subset of the most important promoter (from CNN models) and non-promoter (from Random Forest models) pCREs identified for each of the seven response groups. The most important promoter pCREs from the CNN models were those that when all values were set to absent (i.e. zero) caused the largest decrease in model performance (see **Methods**). The most important pCREs from the Random Forest models are those that when used at a node in a decision tree, were able to best separate genes by their response group (see **Methods**). The importance scores of all pCREs based on these two approaches are in **Table S6, S7**.

We first characterized the multi-omics signatures of the most important promoter pCREs using the additional types of omics information described above, by determining how much more frequently the sites of each promoter pCREs overlapped with each of the additional omics information in response group genes than randomly expected using a set of 1,000 random *6*-mers (**Table S7**). Focusing on the top five most important pCREs from each response group, we found that these pCREs could be clustered into three groups based on their degrees of overlap between their sites and the additional omics information (**Fig. 6A**). Group 1 pCREs were unique in that, in addition to overlapping with known TF binding (DAP-Seq) and chromatin accessible (DHS) regions, they were also much more likely to overlap with conserved noncoding sequences (CNS) than random *6*-mers (dashed boxes; **Fig. 6A**), suggesting these pCREs are more highly conserved across the *Brassicaceae*. Group 2 pCREs also frequently overlapped with DAP-Seq and DHS regions, although to a lesser extent, and were also less likely to overlap histone marks associated with active transcription (e.g. H3K23ac, H3K4me1), which was notable given how many important pCREs identified for the down-regulation response groups (i.e. DNN, DND, NND) clustered into Group 2. Finally, Group 3 pCREs were less likely to overlap with DAP-Seq regions than random *6*-mers, suggesting these pCREs may be bound by TFs not yet included in *in vitro* binding databases.

**Fig. 6.**
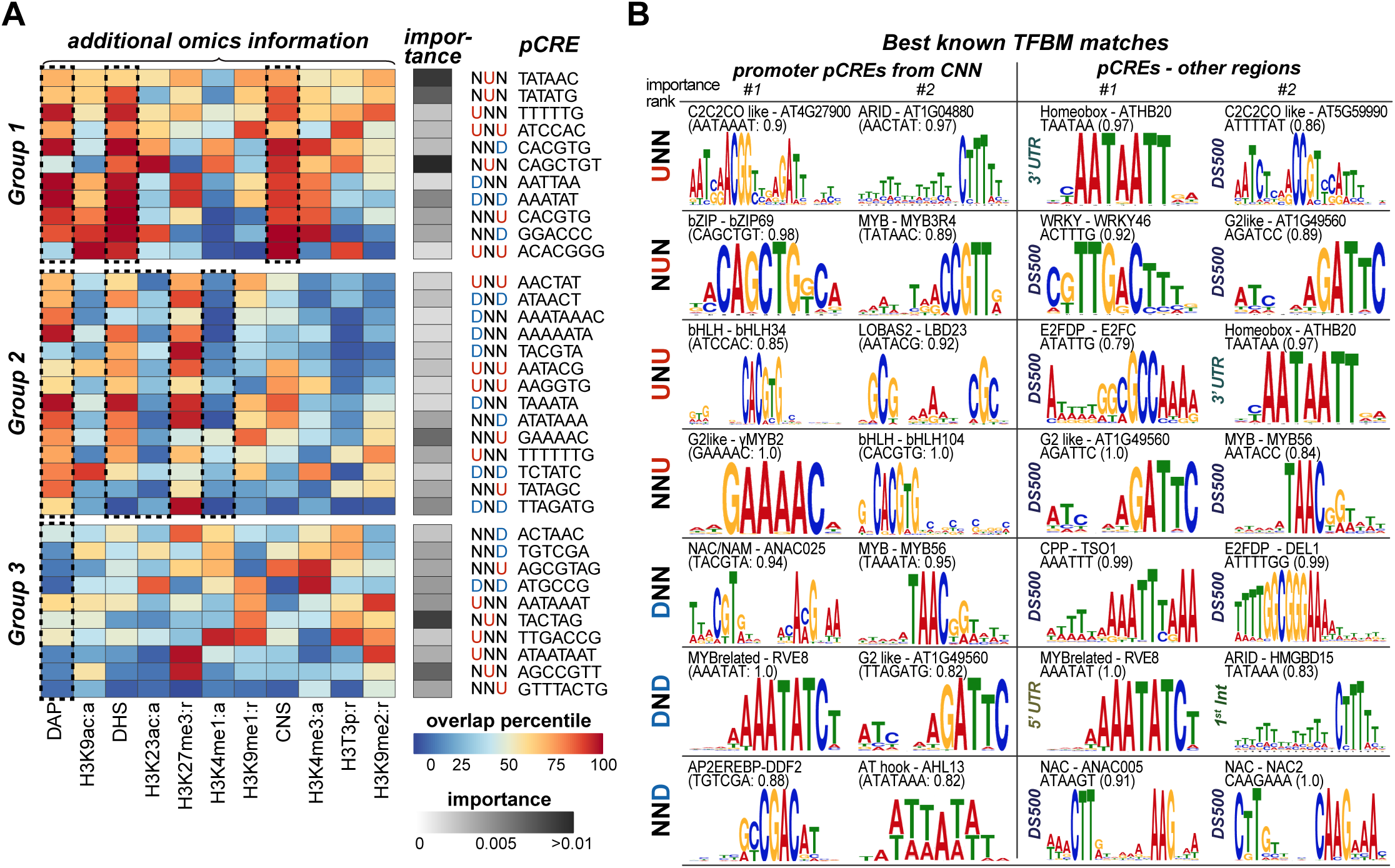
Overview of the most important pCREs for our models of the cis-regulatory codes. (A) The top five most important promoter pCREs from CNN models clustered using k-means clustering (k=3) into three groups based on the pattern of overlap between their sites with additional omics information and sorted using hierarchical clustering. The overlap percentile refers to how frequently a pCRE overlaps with each additional omics information in the promoter of response group genes compared to 1000 random 6-mers, with values in darker red signifying higher degrees of overlap compared to the random background. The importance score is the median decrease in model performance on the test set when a pCRE and its associated additional omics information is removed from the CNN model (i.e. larger decrease in performance means a larger importance). (B) The TF name, motif logo, and sequence similarity score (Pearson’s Correlation Coefficient; PCC) for the known TFBMs that best match the top two promoter pCREs from the CNN model (left two columns) and the top two non-promoter pCREs from the RF model using pCREs from all five gene regions (see purple in Fig. 5A) (right two columns) for each response group.

We next characterized promoter and non-promoter pCREs by determining which were similar to known TFBMs and which represented putatively novel CREs (see **Methods**). Across all pCREs we identified, 40.5% of promoter pCREs and 37.6% of pCREs from other regions were significantly similar to a specific known TFBM (i.e. sequence similarity (PCC) was > 95^th^ percentile of PCCs between TFs in the same family) (**Table S6, S7**). Focusing on the two most important promoter and non-promoter pCREs for each response group (**Fig. 6B**) we found many different TFs and TF families represented. The promoter and non-promoter located pCRE for the DND models, AAATAT, is identical to the TFBM of a MYB related TF, *REVEILLE8* (*RVE8*) (**Figure 6B**), which has been proposed to be involved in a negative feedback loop regulating the circadian clock’s response to temperature (55). The most important non-promoter pCRE for the NUN model, ACTTTG, is similar to TFBMs in the WRKY TF family (PCC to *WRKY46* = 0.92), which are known to be involved in osmotic and salt stress response (73). The most important promoter pCRE for the NND models, TGTCGA, is similar to TFBMs in the AP2 TF family (PCC to *DDF2* = 0.88), which are known to be involved in heat, cold, and drought tolerance in *A. thaliana* (74). Taken together, these three examples give us confidence that our approach to modeling and interpreting the *cis-*regulatory codes is useful because it allowed us to find pCREs similar to known TFBMs for TFs known to be involved in heat, drought, and combined heat and drought stress.

In contrast, the most important pCREs for the NNU response group are not similar to TFBMs for TFs known to be involved in either heat or drought stress. For example, the most important promoter pCRE, GAAAAC is identical to the TFBM for the G2-like *γMYB2* TF, which has no known association with stress response. The second most important promoter pCRE, CACGTG is identical to the TFBM for *bHLH104*, which while known to be involved in regulating iron homeostasis in *A. thaliana* (75), is not associated with other stresses. Similarly, the most important non-promoter pCRE for NNU, AGATTC, is identical to the TFBM for AT1G49560, a G2-like family TF possibly involved in regulating flowering time. This highlights the need for further study on plant response to combined heat and drought stress and provides prime putative regulatory elements and associated TFs for further characterization.

In summary, we found that important promoter pCREs belong to three groups that differed in how frequently the pCREs were associated with additional omics information. We also found that while some of the most important pCREs found by our models of the *cis-*regulatory codes were similar to known TFBMs bound by TFs involved in heat and/or drought stress response, others (i.e. those enriched in NNU genes) were similar to TFs with no established association to either stress condition. Taken together, these findings highlight the complexity of the *cis*-regulatory codes of response to single and combined heat and drought stress in *A. thaliana* and the need for further study.

## DISCUSSION

Understanding how plants regulate their response to combined heat and drought stress is of great importance because of the frequency with which these stresses co-occur and severity of their impact on our agricultural sector (36). Here we identify candidate pCREs and develop models of the *cis-*regulatory codes regulating response to single and combined heat and drought stress in *A. thaliana*. We found that presence/absence of candidate pCREs could predict heat and drought stress transcriptional responses and that incorporating additional omics information (i.e. chromatin accessibility, sequence conservation, known TF binding, and histone markers) and pCREs outside of the proximal promoter region and improved model performance. We also explored the use of a deep learning approach, CNN, to integrate multi-omic input data and demonstrated that it performed better than Random Forest, a classical machine learning algorithm. Further, by interpreting our models of the *cis-*regulatory codes, we were able to provide novel biological insights, including identifying which pCREs and additional omics information were most important for predicting response to single and combined heat and drought stress. These important pCREs are prime targets for follow up characterization.

Because our models are not able to perfectly predict a gene’s response group, there is still more to learn about the complexities of the regulation of response to single and combined heat and drought stress. One factor that is limiting our ability to model the *cis-*regulatory codes is that genes in a response group are not all regulated by the same mechanisms. This issue is compounded by the fact that samples were gathered only at a single time point a few days after the stress conditions were applied. From this snapshot we cannot determine whether the stress responsive genes began to respond immediately after stress initiation or later after the plants began to acclimate, limiting our ability to separate genes with different dynamic responses to combined stress (76). A second limiting factor is that we are missing critical information about the rate of mRNA degradation. Because our picture of differential gene expression comes from measuring and comparing the steady state mRNA levels, we cannot determine if the change in gene expression is due to, for example, increase in production or a decrease in degradation. Finally, while incorporating TF binding, chromatin accessibility, and epigenetic mark data into our models of the *cis-*regulatory codes improved their performance, these data were not ideally suited for this study because they were generated from plants at different developmental stages and under different conditions than those used to generate the transcriptomic data (14). This is an important consideration as TF binding, chromatin accessibility, and epigenetic marks change over the course of development and in response to environmental conditions (45, 77, 78).

The regulatory codes underlying how plants respond to stressful environments involve many molecular players acting in interconnected ways. Stress responses are also dependent on countless other factors such as the duration (79), severity (80), and frequency (81) of the environmental stress and the cell/tissue type (82), developmental stage (83), and genetic background (83–85) of the plant. Thus, to more fully decipher these codes, it will be optimal to have multi-omics data with as many of the molecular players in place as possible, across multiple time points, in a myriad of environmental conditions, at different developmental stages, and from different tissue and cell types. However, access to such a dataset alone will not improve our understanding of plant stress response. Rather, computational approaches that can integrate and find patterns in such heterogeneous data are crucial. Further, the models generated need to be interpretable so that we can derive new biological insights from them. Our study represents one such interpretable modeling approach. Although there is a substantial room for improvement, our general approaches can be used to better understand the regulation of other developmental and stress induced responses in plants and other organisms.

## ACKNOWLEDGMENTS

We thank the many members of the Shiu lab that have provided valuable suggestions to the project, especially Melissa Lehti-Shiu, Sahra Uygun, Nicholas Panchy, Bethany Moore, Siobhan Cusack, Ming Jung Liu, and Peipei Wang.

