## Supplemental Figures for "The *cis*-regulatory codes of response to combined heat and drought stress in *Arabidopsis thaliana*"

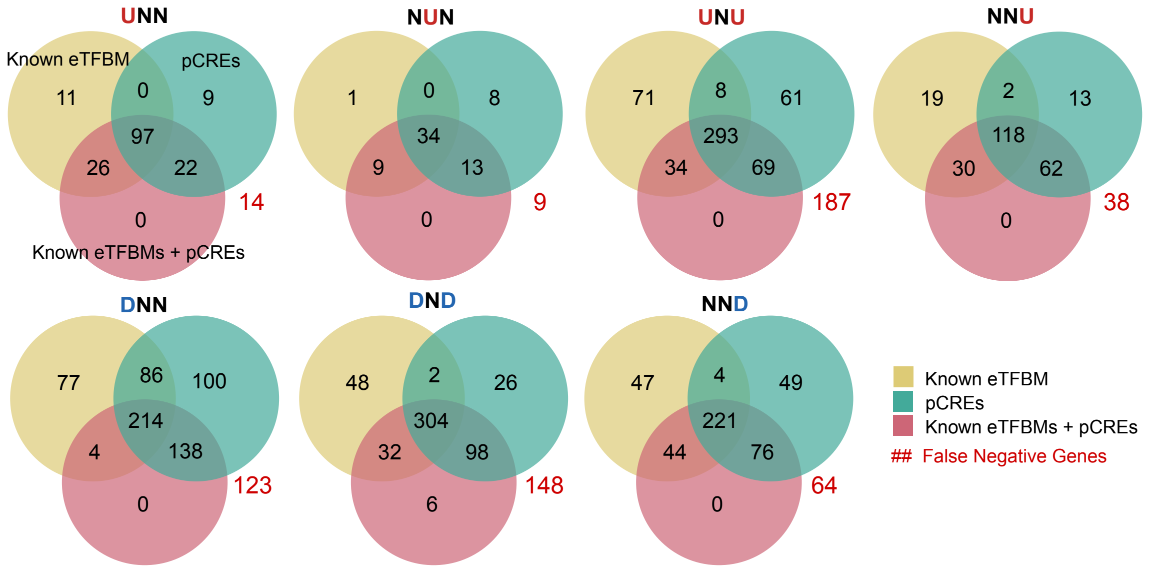


Fig. S1. Overlap in true positive gene predictions from models using known eTFBMs, pCREs, or both features as input.

The number of genes that were similarly correctly predicted by known eTFBM, pCRE, and known eTFBM + pCRE models of the cis-regulatory code for each response group. The number of genes in a response group that were predicted as belonging to that response group by any of the models (i.e. False Negatives) is shown in red.

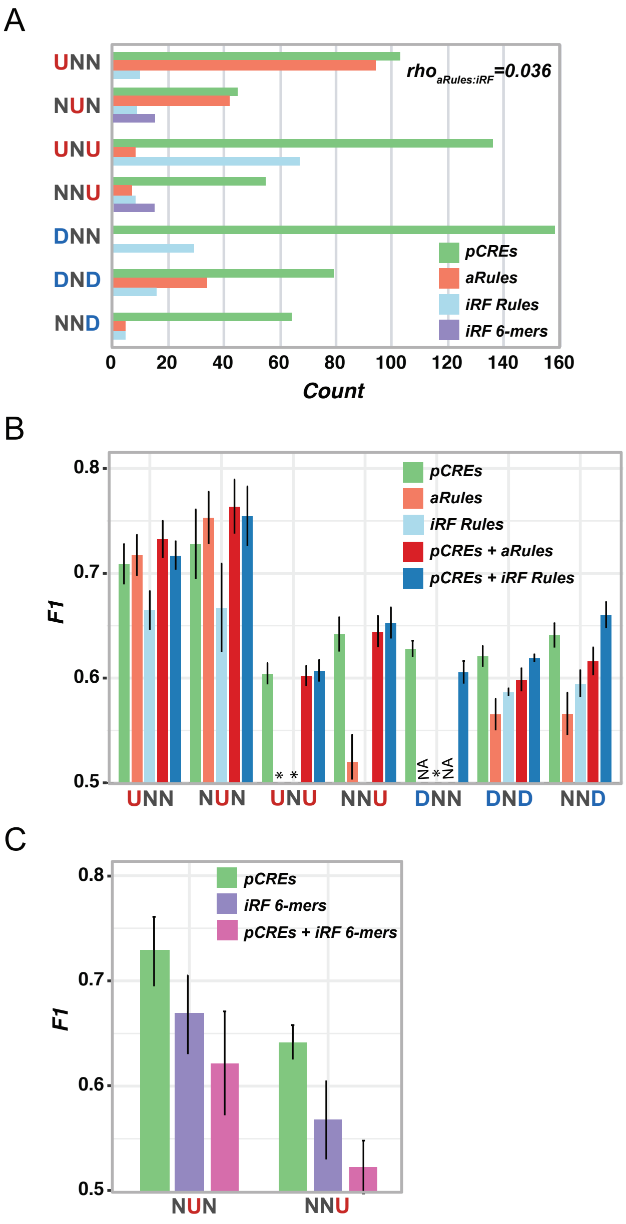


Fig. S2. Impact of including association rules between pCREs as model features.
(A) The number of pCREs (green), and pCRE association rules identified using aRules (pink), and iRF rules (light blue) for each response group (Y-axis). (B) Performance of RF models of the cis-regulatory codes using all single pCREs (green; as in Fig. 3A), only aRules (pink), only iRF rules (light/), single pCREs + aRules (red), and single pCREs + iRF rules (dark blue) as input features. (C) Performance of RF models of the cis-regulatory codes using 6-mer pairs identified by iRF from a set of all possible 6-mers as features.

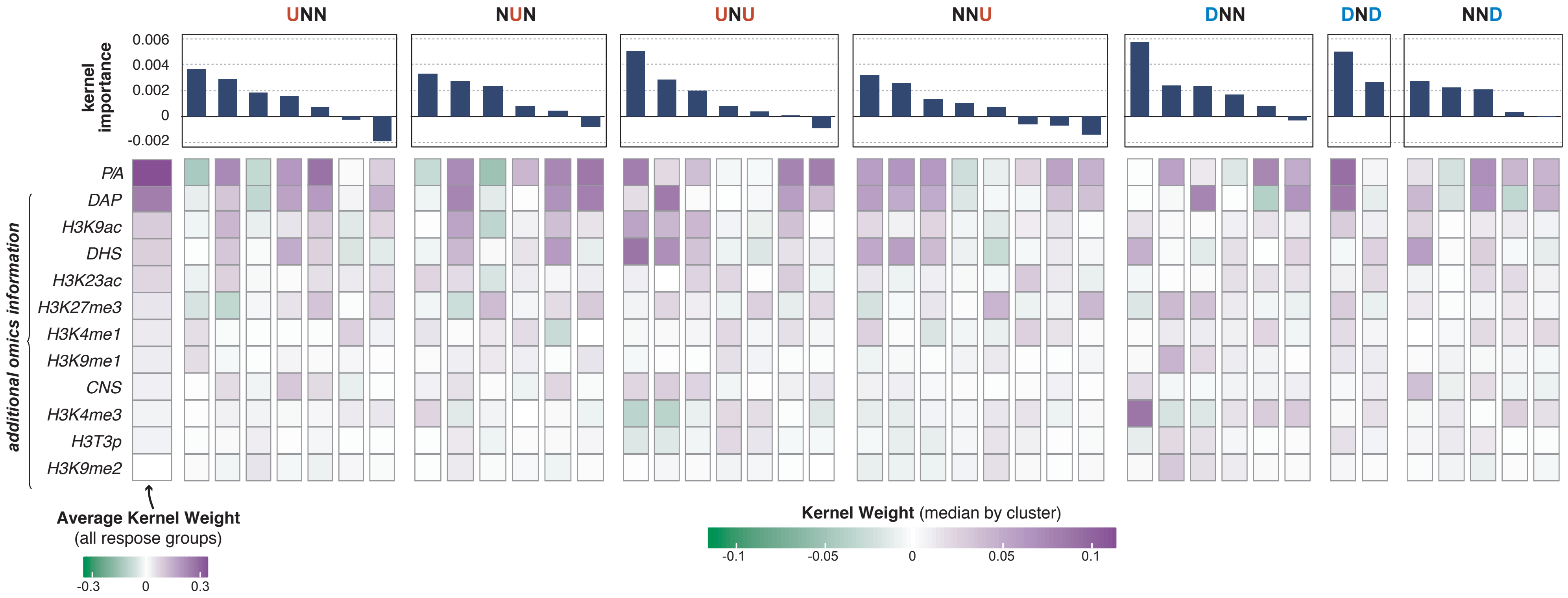


Fig. S3. Probing the trained kernels to understand the important patterns of additional omics information identified by CNN models.
The full results from interpreting the trained CNN models (see Fig. 4C). The feature types (i.e. presence/absence (P/A) and additional omics information) were sorted based on the average kernel weights across all kernels trained for all response groups and replicates (first column). The remaining columns represent kernel clusters for specific response groups. For each response group, all trained kernels from all CNN replicates were clustered using hierarchical clustering with dynamic cutting (min cluster size=250 kernels). The median kernel weights and kernel importance scores are shown here for the resulting clusters.

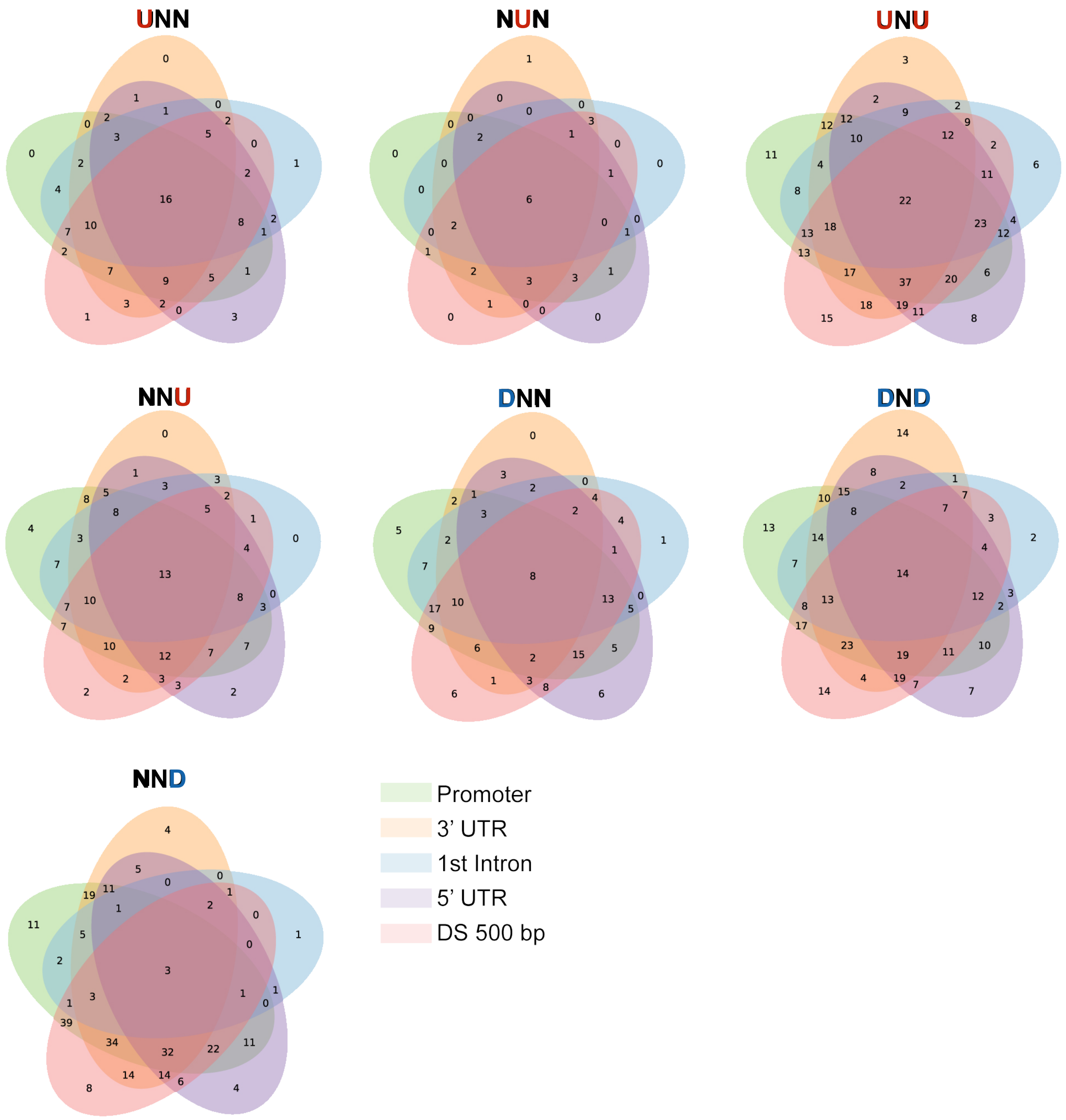


Fig. S4. Overlap in true positive gene predictions from models using pCREs from different genetic regions
The number of genes that were similarly correctly predicted by pCREs identified in the promoter (green), 3' untranslated region (UTR) (orange), first intron (blue), 5' UTR (pur- ple), and downstream (500 bp; DS500; red) regions for each response group.
